# Generic Purpose Pharmacokinetics-Pharmacodynamics Mathematical Model For Nanomedicine Targeted Drug Delivery: Mouse Model

**DOI:** 10.1101/2022.07.13.499855

**Authors:** Teddy Lazebnik, Hanna Weitman, Gal A. Kaminka

## Abstract

Pharmaceutical nanoparticles (NPs) carrying molecular payloads are used for medical purposes such as diagnosis and medical treatment. Currently, the research process of discovering a new applicative candidate for efficient clinical treatment is a time- and resource-consuming process due to the uncertainty of how NP behaves which requires a large number of experiments to study the properties of NPs-based drugs for clinical usage. *In silico* experiments are known to be powerful tools for studying biological and clinical systems and evaluating a drug’s efficiency, which can significantly reduce the number of *in vivo* experiments required. To this extent, in this study, we present a novel spatio-temporal pharmacokinetics-pharmacodynamics (PKPD) model of NPs based drugs. The proposed model takes into consideration the blood flow in the cardiovascular system as well as PKPD dynamics taking place during the drug’s flow and in the target sites. We show that the proposed model has a better fidelity compared to previous models on five *in vivo* experiments with 13 different NPs, done on mice.

## 1 Introduction

The development of nanomedicine-based drugs is a relatively new and promising field in drug discovery [15, 16, 49, 45]. New developments of nanomedicine-based drugs are becoming increasingly more challenging due to the constant increase in costs, decrease in productivity, and attrition of projects as they progress through the development process [24, 55, 39].

One way to handle these issues is using mathematical models and computer simulations to investigate the clinical outcomes of new drugs in a highly controlled manner, relatively fast, and cheap settings as these tools were found to be effective [3, 32]. *In silico* experiments can be used to obtain initial results in the drug development project and thus reduce the number of futile candidates one need to explore *in vivo* [32]. Specifically, Pharmacokinetics-Pharmacodynamics (PKPD) models have been investigated for multiple drugs and environments and shown to well represent the bioclinical dynamics *in vivo* [13, 51, 18].

One approach to model PKPD nanoparticles (NP) based dynamics for drug delivery is using a graph-based model [46]. While the graph-based PKPD model produces a relatively accurate prediction of the biodistribution of the NP-based drug in the body compared to other PKPD mathematical models, it involves several simplifications of the biological systems and the bioclinical properties of the introduced drug giving rise to large errors in the forecast of the corresponding *in vivo* experiments:

- Information about the chemical and physical properties of the NPs is missing, they differ from each other only by their half-life time.
- The blood flow dynamics and their influence on the NPs in the cardiovascular system are poorly modeled. Particularly, it is linear to the amount of NPs in the blood vessels and it depends only on the volumes of the blood vessels.
- The interaction protocol between the NPs and the environment is identical for all the organs and blood vessels.

To simplify some parameters of the elaborated biological systems, invalid assumptions are acceptable in PKPD theoretical models research [61], whereas when using these models as clinical prediction tools to investigate the dynamics of biodistribution, quite large errors are obtained [65]. Therefore, we aim to tackle these inefficiencies as follows: First, we introduce heterogeneous dynamics for each type of NP in the form of interaction and spontaneous protocols that depend on the environment and time. In addition, we take into consideration three chemical and physical properties (size, geometry/shape, and surface material) [75]. Second, we use a single-phase, Newtonian fluid, two-dimensional Navier-Stokes equations [43] with elastic collisions between the NPs and themselves and the blood vessels’ walls. Furthermore, we introduce a heart pulse for the blood flow in a single blood vessel setting and compute the optimal discrete transformation function between two cylinder-shaped blood vessels. Third, we estimate the NP-environment (organ) interaction protocol using a system of ordinary differential equations that are heterogeneous to the NP type and the organ taking part in the interaction as they fitted on multiple organs and NPs simultaneously.

Based on the proposed model, we obtain the contribution of each new component on the accuracy of the *in silico* results compared to the corresponding *in vivo* values on five *in vivo* experiments, together involving 13 types of NPs.

The paper is organized as follows. First, we present an overview of PKPD modeling in general and NP-related models in particular, followed by flow dynamics in blood vessels and ordinary differentiation equations (ODE) based methods for PKPD modeling. Second, we proposed an extended mathematical model for the one proposed by [46], inspired by nanomedicine dynamics. Third, we present the flow dynamics component. Fourth, we show an ODE-based NPs-environment generic PKPD component. Sixth, a simulation based on a mouse’s anatomy for each component independently and the overall model is provided. Seventh, a comparison between the simulation’s (*in silico*) results and the *in vivo* values is provided. Finally, we discuss the main advantages and limitations of the model compared to other PKPD models.

## 2 Related Work

The process of drug discovery and investigation in general and of nanomedicine, in particular, is complex, time-consuming, and requires researchers from various disciplines such as biology, engineering, and chemistry [49, 45]. Nanomedicine is the field of biodegradable and biomimetic nanoparticles (NPs) that are used for multiple clinical procedures. More often than not, NPs are considered objects whose dimensions are in the scale between 1 to 100 nm with a wide range of materials, shapes, payloads, etc [53].

One promising usage of NPs is that they can serve as drug delivery agents by efficiently carrying and delivering drugs to specifically targeted sites and releasing them in a controlled manner. Drugs are loaded on NPs and injected into the bloodstream as nanomedicine therapeutics, designed to modify the pharmacokinetics (PK) and pharmacodynamics (PD) properties of their associated drugs for efficiently targeted drug delivery [54].

*In silico* experiments which are based on robust mathematical models of the biological, physical, chemical, and clinical processes are a powerful tool to develop and investigate nanomedicines [3, 19], as conducted in this research. Specifically, how to develop a robust simulation of PKPD NPs-based drugs *in vivo* dynamics that takes into consideration multiple processes in several levels in order to achieve high accuracy.

A review of the biological properties of NPs-based targeted drug delivery treatment is presented. In which, the versatility of the treatment and its dynamics in different time frames is provided. Afterward, a mathematical description of NPs’ flow in the cardiovascular system is offered, showing several approaches for the flow of particulars in the blood flow. Finally, several PK studies described the usage of such models on *in vivo* data to investigate the dynamics of NPs-based treatments are described.

### 2.1 Targeted drug delivery nanomedicine

Precision medicine is propelled by technologies that enable genomic analysis, molecular profiling, and optimized drug design to better tailor treatments for individuals [52]. Although precision medicine has resulted in some clinical successes, the use of many potential therapeutics has been held back by pharmacological issues, including toxicities and drug resistance [27].

During the last several decades targeted (controlled) drug delivery methods have advanced significantly, leading to various clinical formulations which improve patient compliance and convenience [38]. Recent developments allow delivery of drugs at desired release kinetics for extended periods, substantially improving drug efficacy and minimizing side effects [7]. In particular, NPs made from natural and synthetic materials have received the majority of attention due to their stability and ease of surface modification for targeted drug delivery [36]. They can be tailor-made to achieve both controlled drug release and disease-specific localization by tuning the core characteristics and surface chemistry [58]. It has been established that nanocarriers can become concentrated preferentially to tumors, inflammatory sites, and antigen sampling sites by the enhanced permeability and retention (EPR) effect of the vasculature [68]. Once accumulated at the target site (for example, cancer tumor), NPs can act as a local drug depot, providing a source for a continuous supply of encapsulated therapeutic compounds at the disease site [68].

### 2.2 Blood flow with nanoparticles

Simulation of fluid flow is a well known challenge in general [74, 34, 70] and for blood flow in particular [50, 29, 31]. An general description of the blood flow in a single blood vessel can be obtained by treating the blood as an incompressible, single-phase, Newtonian fluid governed by the Navier–Stokes (NS) equations:

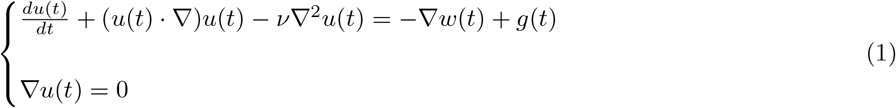

where, *u* denote the velocity of the fluid, 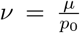 is the kinematic viscosity, *w* is a specific, per unit mass, thermodynamic work, *p* is the pressure of the blood, and *g* is the body acceleration acting on the fluid [63]. By using this version of the NS equations, it is assumed that the blood satisfies Galilean invariant stress and isotropic [40]. While this representation can approximate the average blood flow dynamics, it lacks several components such as spatial dynamics, dynamic geometric configuration (and therefore dynamic, time-depended, boundary condition) of the blood vessel, and general body-related processes such as heartbeat [63].

Liu and Li [50] modeled the blood vessel flow dynamics using a coupling of the NS equations for the inner part of the blood vessel where the blood flow with elastic Navier-Lame equations for the blood vessel’s wall by treating it as a linearly elastic material. The authors approximate the blood vessel’s geometry using a cylinder that is static in time, relaxing the realistic dynamic border condition. At this point, due to the lack of biological data on the elasticity of blood vessels and the effect of the surrounding on them *in vivo* - this approach currently remains theoretical. In our model, we relax the elastic assumption of the blood vessel’s walls, treating it as a static boundary condition for the blood flow dynamics while being elastic for NPs collisions.

As a consequence of the radial symmetry of the cylinder geometric configuration, the NS equations can be reduced from a three-dimensional Cartesian axis system into two-dimensional radial axis system [14]. In such cases, the blood flow NS equations (see Eq. (1)) can be modified to:

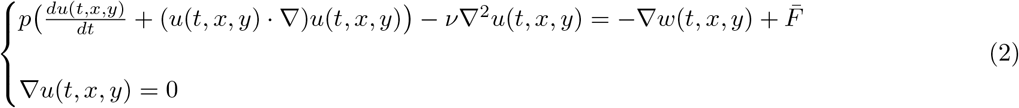

such that *p* is the pressure on the blood and

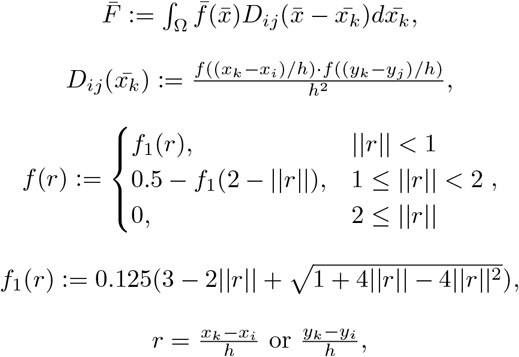

where *h* is the Euclidean grid size and (*x*_*i*_, *y*_*i*_) are the Euclidean points in space [66]. Using this representation, one can numerically compute the radial and axial velocity vector fields of the flow [43, 21].

In addition, Belardinelli and Cavalcanti [10] proposed an extended nonlinear two-dimensional NS model of the blood flow with elastic vessels. Namely, the boundary condition Ω is changing over time due to wall motion and instantaneous taper angle taken into consideration in the model. This allows the calculation of axial and radial velocity profiles with relatively low computational complexity on one hand while making the entire computation less stable in time as a numerical error associated with a fluid-boundary interaction is increasing over time. Therefore, in our model, we treat the blood vessel’s wall statically to obtain a more numerically stable simulation at the cost of a worse approximation of the biological system.

Furthermore, blood flow is taking place in multiple blood vessels with different sizes and pressures, which adds a layer of complexity to the simulation as the transformation from one blood vessel to another, results in a sensitive numerical computation [1]. Chow and Mak [22] extended the two-dimensional NS equations with collapsible cylinders in the case of a parallel flow profile [44]. The authors show that the pressure between the blood vessels can be obtained analytically and therefore are numerically stable [22]. However, this analysis which assumes a parallel flow profile does not handle the case where blood vessels bend or one blood vessel is connected to two blood vessels with some opening angle between them.

Lastly, by adding NPs into the flow, the NPs flow inside the blood flow without changing the flow in the arteries due to their nanometric size [60]. Fullstone et al. [30] proposed an agent-based approach to simulate the flow of NPs in the blood flow in a single blood vessel. The authors first computed the velocity vector field using one-dimensional NS equations and afterward computed the changes in this field due to elastic collision with red blood cells (RBCs) represented using a rigid-body [8]. We remove the RBCs from the flow as they are significant only in blood vessels with a small radius. However, we add the elastic collision properties to the NPs to better represent the flow around the blood vessels’ walls.

### 2.3 Fitting PKPD model on in vivo data for NPs-based treatments

PKPD are commonly used to analytically investigate the interactions between a drug and a biological system [13, 51, 62, 17]. More often than not, researchers conduct an *in vivo* or *in vitro* experiment which produces clinical and/or biological data. Afterward, a bio-mathematical model is developed based on the current biological, chemical, physical, and engineering understanding of the processes involved in the conducted experiment. Finally, the model is fitted to the data of the experiment to specifically reflect the experiment with its unique attributes [41, 67].

The PKPD models used for such analysis can be divided into two main groups: specific-purpose and general-purpose models. The specific-purpose models take into consideration the unique properties of the drug and the interaction of the drug with the body as part of the dynamics. As a result, there is a trade-off between specific-purpose and generic-purpose models as the first optimizes the accuracy and better represents the PKPD dynamics of a specific case while getting worse results for any other case. On the other hand, generic-purpose models are performing worse than specific models for the one case they are built for but perform better for any other case in comparison.

An example of a specific-purpose model is the one proposed by Aboring et al. [2] which is a physiologically-based (PI) PK model described by a system of ODEs for the biodistribution of gold nanoparticles in mice. The authors introduce two equations (Eqs. (S17, S18) [2]) which assume the drug is carried in spherical-shaped NPs. Moreover, the authors assume in the model that the drug’s excretion is performed only in the liver [2] which can be different for other types of NPs. Another example of a specific-purpose model is the PI glucose-insulin-glucagon regulatory model in humans proposed by Schaller et al. [64]. The authors used a glucose-insulin dynamics [20] as the main PK process in the model [64] in type 1 diabetes individuals. For any other type of disease, the model is invalid. Similarly, Tsuchihashi et al. [71] proposed a PKPD model for doxorubicin in long-circulating liposomes in mice. In the model, the authors assume the drug doxorubicin (DOX) has two phases - liposomal DOX and free DOX such that liposomal DOX is converted into free DOX during the treatment. Naturally, any drug that does not fulfill this property can not be modeled by the model proposed by Tsuchihashi et al. [71].

## 3 Biological Model

### 3.1 Inspiration From Nanomedicine Dynamics

Inspired by biological nanomedicine dynamics, one method of clinical treatment via NP-based drugs is as follows. The treatment starts by injecting a population of NPs into the cardiovascular system from a single blood vessel. During the treatment, more NPs can be injected using the same or different blood vessels. The NPs are carried in the bloodstream to different organs and tissues in the body. When NP enters an organ or a tissue it interacts with the local environment thus the interactions are different for each organ and tissue, and with other NPs, in a stochastic manner. Such interactions happen in the bloodstream as well but in a relatively small amount compared to the organs due to the flow dynamics.

An NP is a complex drug carrier from both biological and engineering prospective as it has a large number of properties. Nevertheless, there are six properties that have major direct influence on their dynamics in the body and ability to perform a given task: size [13], shape [51], core material [47], surface material [11], and payload [52]. Both the interactions between an NP and the environment and between the NP and other NPs may influence the NP (for example, release its payload / modify its shape) and therefore the way the NP float in the cardiovascular system and how it interact with its environment (organs) [2].

### 3.2 Model Definition

Inspired by the dynamics of the biological systems (Section 3.1), we propose a mathematical model that represents the behavior of targeted drug delivery NP-based drugs within a biological system. The model is defined by a tuple *M*(*t*) := (*D*(*t*), *E*(*t*)) where *D*(*t*) is a set (population/swarm) of the medical NPs (agents) and *E*(*t*) is the environment which describing the geometry (cardiovascular system and organs) that the agents are located and the biological properties define the possible interaction between the NPs’ and a location, at any given time *t*. The components of the tuple are described below in detail. A schematic view of the model’s structure is presented in Fig. 1.

**Figure 1:**
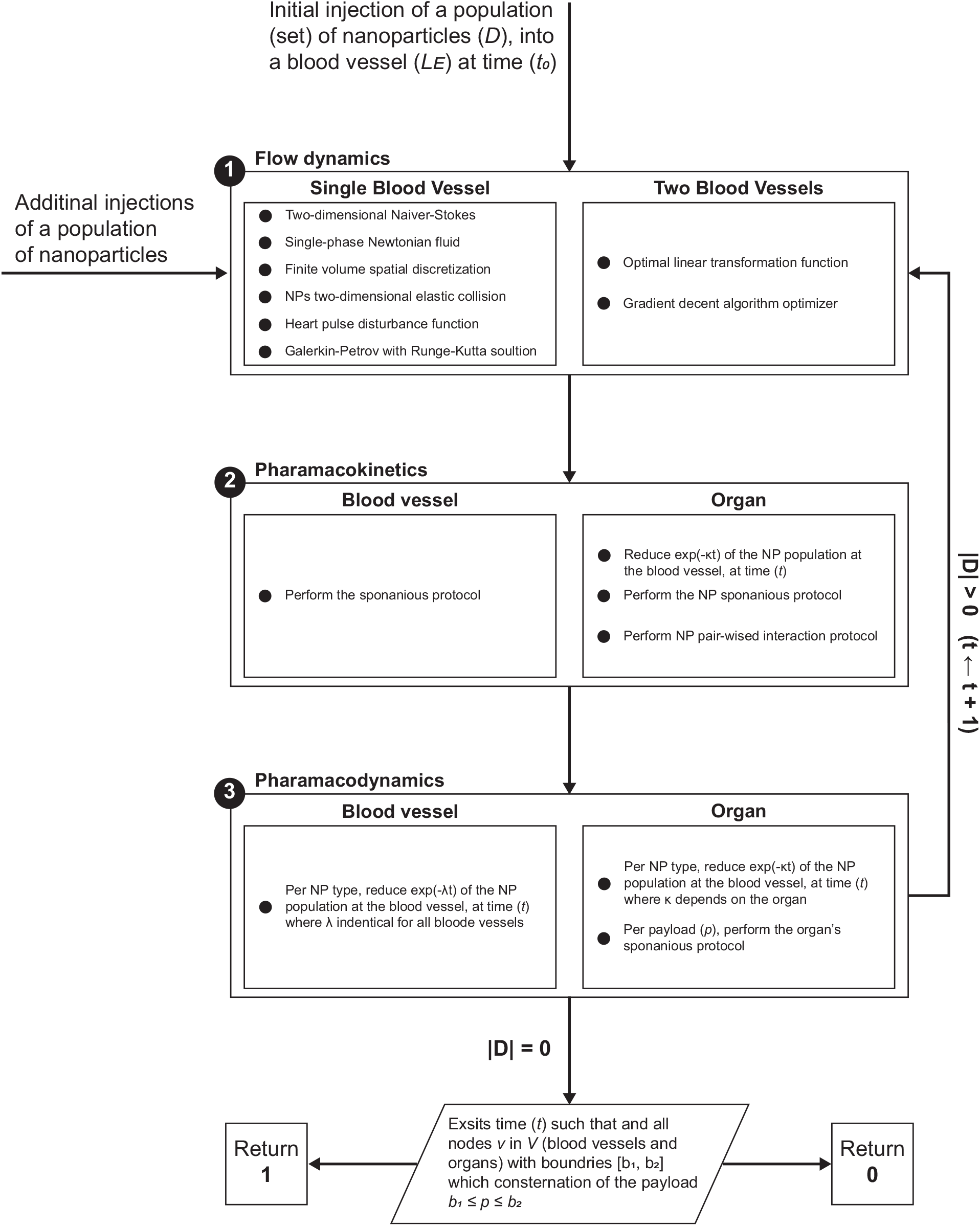
A schematic view of the model’s structure, divided into the three sub-categories of computation with the interactions between them - flow dynamics, pharmacokinetics, and pharmacodynamics.

#### 3.2.1 Nanoparticles swarm drug *D*

The drug in the system is carried by a swarm of medical NPs and therefore it is represented by them. The swarm (*D*) is a set of NP. Each NP in the swarm is defined by an anonymous (unidentified) finite state timed automata with five attributes, two clocks, and two interaction protocols that represented by the tuple: *np* := (*c*_*g*_, *c*_*s*_, *s, g, m, I*_*s*_, *I*_*e*_) where *c*_*g*_ is a global inner clock counting the time pass from the NP’s creation, *c*_*s*_ is an inner clock counting the time pass from the last changes in the NP’s state, *s* ∈ [*s*_*min*_, *s*_*max*_] is the size of the NP, *g* ∈ *G* is the geometry (shape) of the NP where *G* is a set of all possibles NP’s geometries, *m* ∈ *M*_*s*_ is the surface material of the NP where *M*_*s*_ is the set of all possibles NP’s surface material, *I*_*s*_ is the spontaneous dynamics of the NP, and *I*_*e*_ is the interaction dynamics between the NP and other NPs. The size attribute is continuous rather than discrete. Therefore, we assume there is a value Δ*s* such that two sizes *s*_1_, *s*_2_ are inseparable iff |*s*_1_ − *s*_2_| ≤ Δ*s*. As a result, it is possible to represent the size attribute discreetly by dividing the range into Δ*s* bins.

The spontaneous dynamics of an NP (*I*_*s*_) is the biological and chemical processes that change one or more of the NP’s attributes such that it is independent of the possible interactions of the NP with other NPs and drugs but may be time-dependent. Therefore, *I*_*s*_ is defined by a function *I*_*s*_: ℝ^2^ × ℕ^5^ × *L*_*E*_ → ℕ^4^ × *L*_*E*_ such that *I*_*s*_(*c*_*g*_, *c*_*s*_, *s, g, m, a, l*) → (*s, g, m, a, l*) where *l* is a LE (*L*_*E*_ ∈ *E*). For example, after time *T* from the creation of the NP, the payload drug becomes inactive and therefore *I*_*e*_ modifies the payload (*p*) attribute when *c*_*g*_ *> T* for the first time.

The interaction dynamics of a NP (*I*_*e*_) is the biological and chemical processes that change one or more of the NP’s attributes due to interaction with the other NP (*np* ∈ *D*) in the context of the environment *E* where they are located. Therefore, *I*_*e*_ is defined by a function *I*_*e*_ : *np* × *np* × *L*_*E*_ × ℝ → *np* × *np* × *E* such that *I*_*e*_(*np*_1_, *np*_2_, *L*_*E*_, *t*) → (*np*_1_, *np*_2_, *L*_*E*_), where *np*_1_ and *np*_2_ are two NPs (*np*_1_ /= *np*_2_), *L*_*E*_ is a LE (*L*_*E*_ ∈ *E*) such that *np*_1_ and *np*_2_ located at *L*_*E*_ ∈ *E* due to the interaction graph (*FG*), and *t* is the time pass from the beginning of the dynamics.

##### 3.2.2 Environment *E*

The body can be treated as one environment (*E*) which divided into several local environments (*L*_*E*_) such that ⋃*L*_*E*_ = *E* and 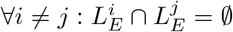, where each one has unique properties that has a different effect on NPs. For example, the liver may deactivate an NP with a highly toxic payload while ignoring other NPs. A LE *L*_*E*_ is defined by the way it interacts with other NPs, its chemical and biological processes which are independent of the NP, and its chemical state at a time *t*. Therefore, we define *L*_*E*_ as a tuple *L*_*E*_(*t*) := (*l, I*_*s*_(*t*), *d*(*t*), *o, g*(*t*)) such that *l* is the indicator of the LE used by the interaction dynamics (*I*_*e*_) of the NPs, *I*_*s*_ is the spontaneous dynamics of the LE which modifies *d* independently to the interactions with the NPs as a function of time, *d* is an injective map function *d* : 𝔻^*n*^ → ℝ^*n*^ where 𝔻 is the set of all drugs *drug*_0_, …, *drug*_*n*−1_ in the system mapped to the amount of each drug in the LE, *o* is a binary property that marks if the LE is a part of the cardiovascular system or not, and *g* is the geometrical configuration of the LE which is defined by triangulation of the surface over time.

The spontaneous dynamics of a LE (*I*_*s*_) are the biological and chemical processes that change *d* which are not a result of the interactions with the NPs. Therefore, *I*_*s*_ is defined by a function *I*_*s*_ : 𝔻^*n*^ × ℝ → 𝔻^*n*^ such that *I*_*s*_(*d, t*) → *d*. For example, in the liver, any drug with high enough toxicity is evacuated from the body over time while others are kept unchanged.

In addition, a flow graph *FG* is defined as a directed, non-empty, connected graph where the nodes are the LEs in *E* and the edges are the abstract connections between the physical nodes.

##### 3.2.3 Scheduler

The model has a synchronized clock. In each clock tick (marked by *t*_*i*_ for the *i*_*th*_ tic) the following three actions occur:

1. The interaction graph *FG* is updated according to the mobility of the NPs’ population (*D*) in the environment *E*.
2. For each NP in the population, in a random order, the spontaneous dynamics (*I*_*s*_) and pair-wise interaction dynamics with other NP (which is a neighbor of the first NP and does not take part in interaction dynamics in the same clock tick) are performed.
3. For each local environment *L*_*E*_ ∈ *E*, in a random order, the spontaneous dynamics function (*I*_*s*_) is performed.

### 3.3 Drug Delivery Successful Treatment

In the case of drug delivery NPs based treatment, a treatment is considered successful if it fulfills two conditions: a given concentration of drugs reached a given set of target LEs (*T*_*E*_) over the course of the treatment, and the non-target LEs (*E* \ *T*_*E*_) keep a given concentration of drugs (not necessarily equal) over the course of the treatment. For example, assume we wish to treat cancer by injecting NPs that statistically penetrate to cancer cells and not normal cells. Such treatment was considered successful if enough NPs penetrated the cancer cells to kill them on the one hand but did not penetrate to normal cells too much and by that killing the patient. One can formally define a *successful* treatment as follows:

#### Definition 3.1.

Given the set of concentrations 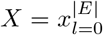 such that each *x* ∈ *X* : 𝔻^*n*^ × ℝ^2*n*^ → {0, 1} for each *L*_*E*_ ∈ *E*. A treatment is defined *successful* iff ∃*t* : (|*D*| = 0 ∧ ∀*x* ∈ *X* : *x*(*t*) = 1).

## 4 Flow Dynamics

We implement the mobility of the swarm of NPs (*D*) inside a single blood vessel as the local environment (*L*_*E*_) and in the transformation between two blood vessels (*N* (*L*_*E*_)) using the agent-based simulation approach under blood flow [30].

### 4.1 Model Definition

The flow dynamics model is a tuple *F*_*d*_ := (*D, L*_*E*_, *N* (*L*_*E*_)) such that *D* is the population of NP located in a local environment *L*_*E*_ during the flow dynamics, *L*_*E*_ is the local environment where the flow dynamics takes place, and *N* (*L*_*E*_) ∈ *E* are all the nodes in the environment (*E*) that are neighbors of *L*_*E*_. The components of the tuple are described below in detail.

The NPs swarm, donated by *D*, is extending the definition provided in Section 3.2.1 by adding four parameters: two-dimensional acceleration (*a*), velocity (*v*), and location (*x*) vectors, and mass (*m*). The vectors are represented in a radial coordinate system (e.i. *r, h* where *r* is the distance from the center of the geometry and *h* is the distance from the initial surface where NPs are introduced into the geometry). These parameters define the movement of the NPs inside the blood vessel. The mass of the NP is obtained using:

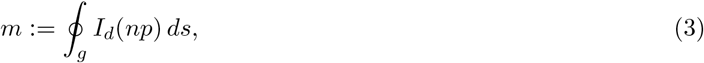

where *I*_*d*_ is a predicate function that for each three-dimensional segment *s* ⊂ *g* computes the weighted average mass of *np* in the segment. Therefore, the movement of a NP in each clock tic (*t*_*i*_) is defined using the formula:

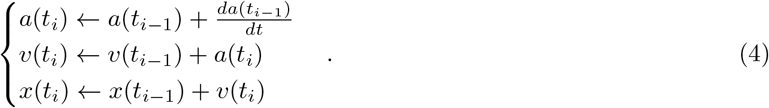

The local environment *L*_*E*_ is extending the definition provided in Section 3.2.2 by adding three parameters and a cylinder geometrical configuration: 1) the radius of the blood vessel *R* is assumed to be constant through the blood vessel and in time; 2) the length of the blood vessel *H*; and 3) the average pressure **P** of the blood over time.

### 4.2 Single Blood Vessel Flow Dynamics

At each clock tick, according to the model’s scheduler (see Section 3.2.3) the swarm of NPs (*D*) flows in the environment. The following three biological and physical processes take place as part of the NPs’ flow in a single blood vessel. These dynamics are referred to veins and arteries as the physical and biological processes in capillaries are significantly different. First, the acceleration vectors of the NPs *D* are modified by the vector field that is defined by solving two-dimensional NS equations [43, 21]. Second, a heartbeat adds distortion to the velocity field of the blood in the blood vessel. Third, a two-dimensional elastic collision between one NP and another and between an NP and the blood vessel’s wall is modifying the velocity (*v*(*t*_*i*_)) and the location (*x*(*t*_*i*_)) of the NPs.

#### 4.2.1 Two-dimensional Navier-Stokes

At each step in time (schedule’s tic), the flow dynamics (e.g., Eq. (1)) performed on the blood vessels. The order is dictated by the distance of the node from the *heart* node in a topological sort [59] of the flow graph. At each blood vessel, the NS is treated as a single-phase Newtonian fluid with finite-volume radial-height discretization.

##### Single-phase Newtonian fluid

In the simulation, we assume the blood is a single-phase Newtonian fluid (SPNF). In physics, an SPNF is defined as follows. First, a fluid is a substance that continually deforms under applied shear stress or external force with zero shear modulus. Specifically, a Newtonian fluid is a fluid in which the viscous stresses arising from its flow, at every point, are linear to the local strain rate [9]. A single-phase fluid is a fluid with the same chemical and physical properties over both space and time. Therefore approximating the blood as an SPNF is allowing to use of a relatively stable version of the NS equations while having a fair approximation of the blood as the mouse’s blood is composed of around 50% plasma and 50% red blood cells in volume [4].

##### Finite-volume radial-height discretization

In the simulation we used a finite-volume radial-height (FVRH) discretization which is illustrated in Fig. 2. Formally, we divide the geometry into sub-spaces such that their volumes are equal. Namely,

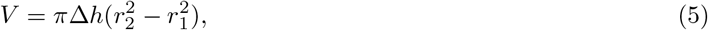

such that Δ*h* is the depth of the segment and *r*_1_, *r*_2_ are the inner and outer radius of a ring circle. We divide each blood vessel depth into 1000 layers, which results in Δ*h* := *H/*1000. Afterward, the radius is obtained by finding *r*_2_ that fulfils Eq. (5) where the first *r*_1_ = 0. The advantage of this discretization over other methods such as finite elements and finite-difference is two-fold. First, radial-height representation is a native oordinate system in a cylinder geometric configuration as used for the blood vessel. Furthermore, as most of the movement is perpendicular to the cylinder’s base and mostly changes due to the distance from the radial center of the cylinder. These biophysical properties make the FVRH discretization a native choice. Second, most of the numerical errors in Eq. (9) and Eq. (6) takes place where *r* is close to 0 and *R*, respectively. On the other hand, the depth inside the blood vessel plays a minor role in these computations. Therefore, by numerically comparing the finite-distances, finite-elements, and finite-volume methods, the last produces the least cumulative error over five blood vessels for 10000 steps in time.

**Figure 2:**
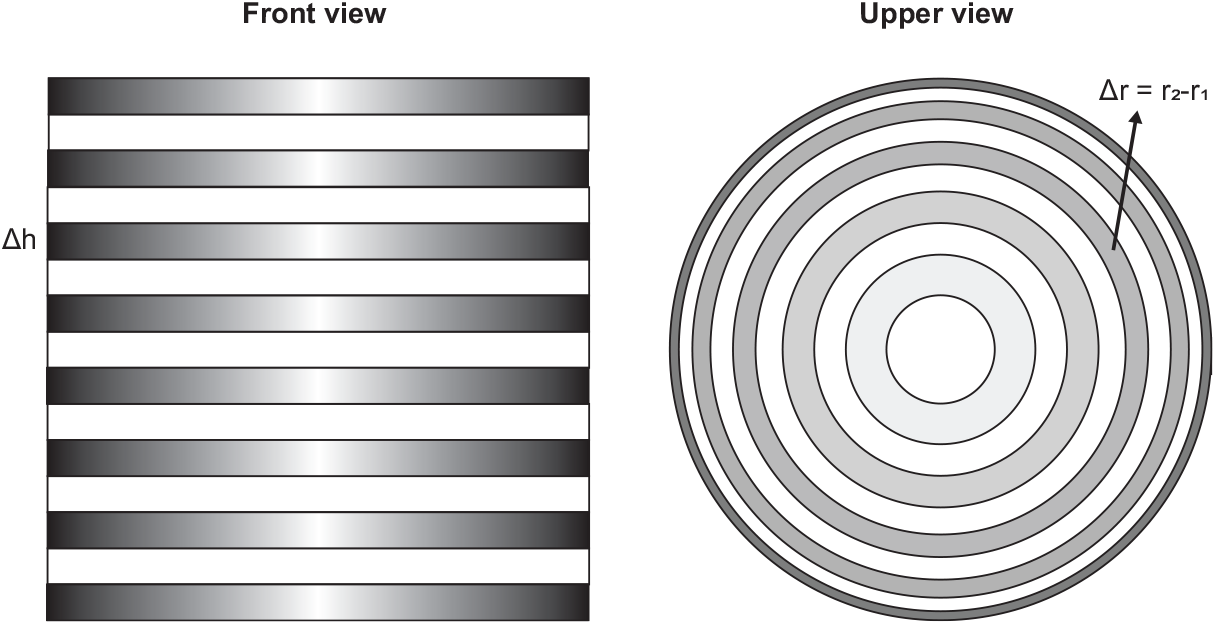
An illustration to the volume-based radial-height discretization. Darker shade means higher rate of sampling (complete white is shown to separate between the components).

### 4.2.2 Heartbeat

We define a heartbeat using a function *pulse* : *t* → ℝ which is introduced on top of the flow dynamics computed by the NS equations (see Section (4.2.1)) operating as a discrete distortion to the pressure of the blood vessel.

The function is formally defined as follows:

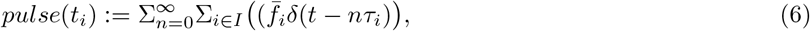

where *δ*(*x*) is the *n*-dimensional Dirac’s delta-function [25] at time *t* − *nτ*_*i*_ and *I* is a set of pulses describing the rhythm of the heartbeat at rest.

#### 4.2.3 NPs two-dimensional elastic collision

We assume NPs have a stiff serve that does not observe any energy from collisions, resulting in a fully elastic collision. Therefore, if any two NPs collide, Eq. (7) describe the changes in the NPs’ velocity [23]:

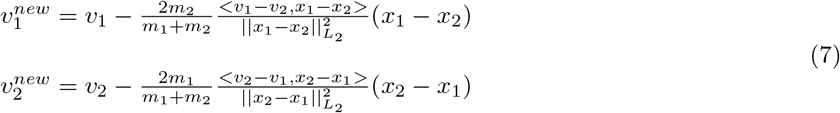

where *x*_1_, *x*_2_, *m*_1_, *m*_2_, *v*_1_, *v*_2_, and 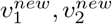 are the locations, masses, the original velocities, and updated velocities of the first and second NPs, respectively.

An elastic collision is performed for each two NPs (*np*_1_, *np*_2_) if exists *τ* ∈ [0, 1] such that

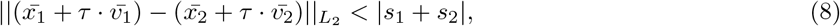

where *s*_1_, *s*_2_ are the sizes of the (*np*_1_, *np*_2_) NPs, respectively. Furthermore, an elastic collision is performed for each NP (*np*) such that exists *τ* ∈ [0, 1] and *h* ∈ [0, *H*] in which

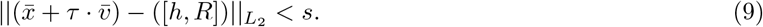

#### 4.2.4 Numerical solution

The two-dimensional NS equations (see Eq. (1)) with the heart beat (see Eq. (6)) are solved using the Galerkin-Petrov (GP) approximation method [69]:

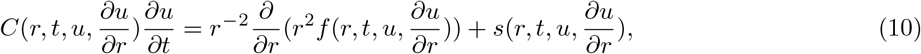

where *m* = 2 and with the finite volume discretization method [42]. In order to simplify the computation, we assume radial symmetrical dynamics (e.g., for any 0 ≤ *r* ≤ *R* the dynamics are equal) in addition to symmetry in 0 ≤ *h* ≤ *H* as the depth of the flow inside the blood vessel has a neglectable effect [33, 48, 43, 6].

In addition, to reduce the error of the approximation and avoid local large errors due to edge cases we compute additional approximations of the 2d NS equations (see Eq. (1)) using the Runge–Kutta–Nystrom method [26, 28], similarly to the GP Eq. (10) method. After obtaining both approximations, we compute the *L*_2_ distance between both resulting vector fields (mark 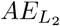). If 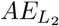 is less than a manually pre-defined threshold *ζ* the average between the two methods is taken. Otherwise, the approximation that closer (e.g., the *L*_2_ distance between the state in time *t* + 1 and *t* is smaller) is chosen. This usage of this process is two-fold. First, it reduces sharp errors in one approximation method by averaging with the second method in general [12]. Second, in extreme cases, by taking the approximation with less change, one may introduce an approximation to the system but in return make it more stable [12].

Afterwards, an elastic collision is performed for each two NPs and between a NP and the blood vessel’s wall if Eqs. (8) or (9) satisfied, respectively.

### 4.3 Transformation between blood vessels

The transformation between two blood vessels is performed in a desecrate manner. Given two circles Ω_1_, Ω_2_ such that defined by the tuple Ω := (*R, f*) where *R* is the radius of the circle and *f* is the velocity vector field vertical to Ω defined for each 0 ≤ *r* ≤ *R* in Ω. A transformation function *T* : (*f*, [0, *R*_1_], [0, *R*_2_]) → ([0, *R*_2_]) between Ω_1_ and Ω_2_ minimize the commutative error over time [57]:

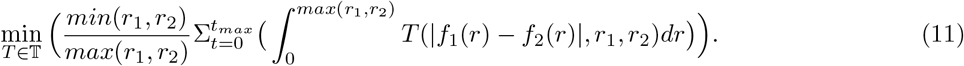

It is possible to solve Eq. (11) for any given *r*_1_, *r*_2_, *f*_1_, and *f*_2_ by define *T* as follows:

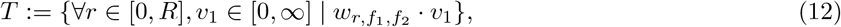

where 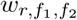 is a weight vector of the transformation *T* for a specific combination of *r, f*_1_, and *f*_2_. In order to find the optimal weights for each *r, f*_1_, *f*_2_ under the constraints that the total error in velocity resulted from *T* is minimal where error is defined as the *L*_2_ distance between the sampled velocity vector from in-vivo experiment and the one obtained from *T*. Therefore, one can find the optimal *T* using the gradient descent algorithm over the parameter space 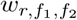 ⊂ *W* [35]. A schematic view of the obtained function *T* is shown in Fig 3.

**Figure 3:**
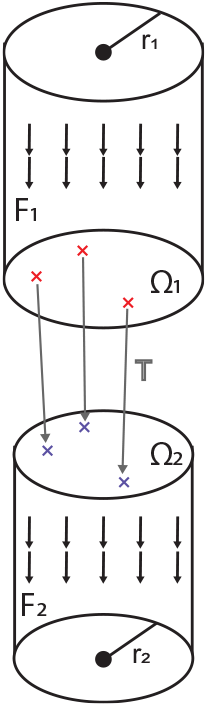
A schematic view of the transformation function *T* between two blood vessels with radius *r*_1_, *r*_2_ which define geometry Ω_1_, Ω_2_ and velocity vector flows *f*_1_, *f*_2_.

## 5 Pharmacokinetics dynamics

It is possible to describe the dynamics of the oncentration of a NP-based drug in the body (*C*(*t*)) as a system of ordinary differential equations each one describing the oncentration of the drug in an organ (*C*_*o*_) and additional one of the circulatory system (*C*_*blood*_) such that *C* = Σ_*o*∈*O*_*C*_*o*_ + *C*_*blood*_. In addition, the drug is a reagent with the immune system (*A*(*t*)) and activates it in some manner. At each step in time, the NP-based drug moves to organs from the circulatory system and the other way around. These processes are generic to any type of NP (and between the different states of nanorobots) but differ in their parameters. Furthermore, as several drugs may be injected into the body in parallel, the oncentration of all drugs differs by the chemical interactions between them in the context of the organ they are located at.

In Eq. (13), the 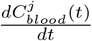 is the dynamical amount of NPs from type *j* ∈ *J* in the circulatory system over time. It is affected by the following three terms. First, each drug *j* has a natural exponential decay in the blood with a half-life of 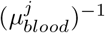. Second, the drug of type *j* reagent immune system response in the blood at a rate 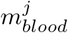. Third, due to a flow dynamics *f*, each drug of type *j* flows from the blood to an organ *o* and back, for every organ (*o*) in the body (*O*).

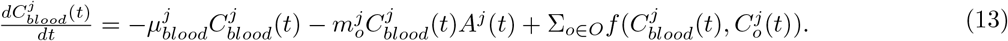

In Eq. (14), the 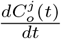 is the dynamical amount of NPs from type *j ∈ J* in the an organ *o* over time. It is affected by the following four terms. First, each drug *j* has a natural exponential decay in each organ *o* with half life of 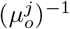. Second, the drug of type *j* reagent immune system response at a rate 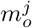. Third, due to flow dynamics *f* each drug of type *j* flows from the blood to an organ *o* and back.

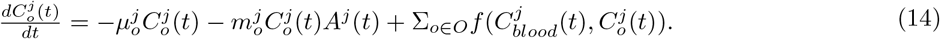

In Eq. (15), the 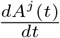 is the dynamical amount of immune system response to the *j* drug over time. It is affected by the following two terms. First, a drug of type *j* ∈ *J* reagent immune system response at a rate 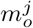. Second, the immune system has a natural exponential decay in each organ *o* with respect to a drug *j* with half life of 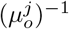.

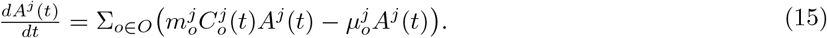

## 6 Model Validation

Based on the proposed model, we examine the performance of the simulation on five *in vivo* studies of NP biodistribution, including 13 types of NPs in total. We used the mouse topology proposed by [46]. The models are divided into four - *baseline, improved flow, improved PK*, and *combined*. The *baseline* model is set to be the model proposed by [46]. The *improved flow* model extends the baseline model by introducing the flow dynamics (see Section 4.2.1). The *improved PK* model extends the baseline model by introducing the generic PK dynamics (see Section 5). The *combined* model extends the baseline model by introducing both the flow dynamics and generic PK dynamics. Furthermore, for each one of the experiments the NP’s population, injection node, and the biological properties taken into consideration would be modified to fit the *in vivo* settings.

The simulation (Algo. 1 in [46]) with the proposed extensions (see Sections 3-5) was executed rapidly for each type of NP. A population of 1, 000, 000 NPs of the same type ℙ was used for the simulation each time. The population size of 1, 000, 000 is picked since in the flow, the amount of NPs has a direct influence on the biodistribution of the population as a result of the flow dynamics and NPs-organs interaction that both depend on the original size of the NPs population. As a result, we wish to inject a population size as close to the one used in the *in vivo* experiment as possible on one hand. On the other hand, a larger population increases the computation time. To compute multiple tests for each experiment, size of 1, 000, 000 rather than ∼ 150, 000, 000 was picked to balance between these two options. There are 13 NPs that have been used in the simulation as shown in Table 2.

**Table 1:** The pharmaceutical dynamics model’s parameters.

| Parameter | Description |
| --- | --- |
| $\mu_o^j$ | The half life $(\mu_o^j)^{-1}$ of a drug $j$ in organ $o$. |
| $m_o^j$ | The reagent factor of a drug $j$ in an organ $o$ to the immune system. |
| $\mu_j^A$ | The half life $(\mu_o^A)^{-1}$ of immune system cells responding to drug from type $j$ |

**Table 2:** The NP used in the experiment, defined by their size, geometry, and surface material.

| NP | In vivo experiment | Size (nm) | Geometry | Surface Material |
| --- | --- | --- | --- | --- |
| 1 | Ben Akiva et al. [11] | 200 | Spherical | PLGA |
| 2 | Ben Akiva et al. [11] | 220 | Ellipsoid (1d) | PLGA |
| 3 | Ben Akiva et al. [11] | 240 | Ellipsoid (2d) | PLGA |
| 4 | Ben Akiva et al. [11] | 240 | Spherical | RBC |
| 5 | Ben Akiva et al. [11] | 260 | Ellipsoid (1d) | RBC |
| 6 | Ben Akiva et al. [11] | 280 | Ellipsoid (2d) | RBC |
| 7 | Lee et al. [47] | 7 | Spherical | PEGylated-chitosan |
| 8 | Lee et al. [47] | 7 | Spherical | PEGylated-chitosan |
| 9 | Verma et al. [73] | 220 | Spherical | Chitosan |
| 10 | Niu et al. [56] | 132 | Spherical | PLGA |
| 11 | Zhang et al. [76] | 223 | Spherical | P(HB-HO) |
| 12 | Zhang et al. [76] | 241 | Spherical | FA-PEG-P(HB-HO) |
| 13 | Zhang et al. [76] | 241 | Spherical | FA-PEG-P(HB-HO) |

### 6.1 Flow dynamics component

We base the mouse environment topology on the one proposed by [46]. Unlike [46], blood vessels are defined by one node corresponding to the length of its blood vessel.

We conducted three experiments to test the flow dynamics: single blood vessel flow, two blood vessel flow, and single blood vessel flow into two blood vessels flow. In all cases, an initial population of 10000 NPs were equally distributed in the beginning (defined by the direction of the blood flow) of the first blood vessel. In addition, the proposed flow dynamics performance was compared to the flow dynamics proposed by Lazebnik et al. [46] (Eq. (1)) and by Updegrove et al. [72] - named SimVascular - which is assumed to be accurately representing the *in vivo* results and therefore with a fidelity of 1 for any point time.

For the single blood vessel flow, we treat the blood vessels as a torus rather than a cylinder such that an NP that exits the blood vessel’s geometry from the end of the blood vessel allocated back to the beginning of the blood vessel using the function *np*.*h* ← *np*.*h* − *H* where *H* is the length of the blood vessel. The physical and biological parameter values used in Eq. (1) taken from [72].

For the two blood vessel flow, we treat the blood vessels as a torus rather than a cylinder such that an NP that exits the second blood vessel’s geometry from the end of the second blood vessel allocated back to the beginning of the first blood vessel using the function *np*.*h* ← *np*.*h* − *H*_1_ − *H*_2_ where *H*_1_, *H*_2_ are the lengths of the two blood vessels, respectively. For the SimVascular model, the two blood vessels are stitched together such that in the middle there is a cylinder with length (*H*_1_ + *H*_2_)*/*2 and a diameter that linearly increase (decrease) from *R*_1_ to *R*_2_ where *R*_1_, *R*_2_ are the radius of the first and second blood vessels. The second blood vessel is picked to be one of the neighbors of the first blood vessel according to the topology from [46].

For the single blood vessel into two blood vessels flow experiment, we repeated the same setup as the two blood vessel flow but this time for the SimVascular model, the first and each of the two following blood vessels are stitched together such that the angle between the two latter blood vessels forms an angle *α* on the y-axis plan (in a three-dimensional *xyz* model).

Since the experiments were conducted on a mouse cardiovascular topology and biological properties, the parameters of the NS equations were taken according to [72] for the mouse model. In addition, the heartbeat function set to be one single (e.g. |*I*| = 1) with 580 ± 18 beats per minute [37]. Furthermore, the transformation between blood vessels’ function *T* has been obtained using data from the corresponding SimVascular [72] model. The model’s fidelity is defined as the slope of a linear regression between the NPs’ locations according to the tested model and the NPs’ locations of the SimVascular model [46].

The experiment conducted on five blood vessels (Ascending aorta, Right external carotid, Left common carotid, Abdominal aorta G, Abdominal aorta B) from the 84 blood vessels of a mouse in the topology [5], picked manually such that the blood vessels’ length and diameter maximally differ. The results are shown as an average ± standard deviation. A schematic view of the geometrical configurations of the three experiments is shown in Fig. 4.

**Figure 4:**
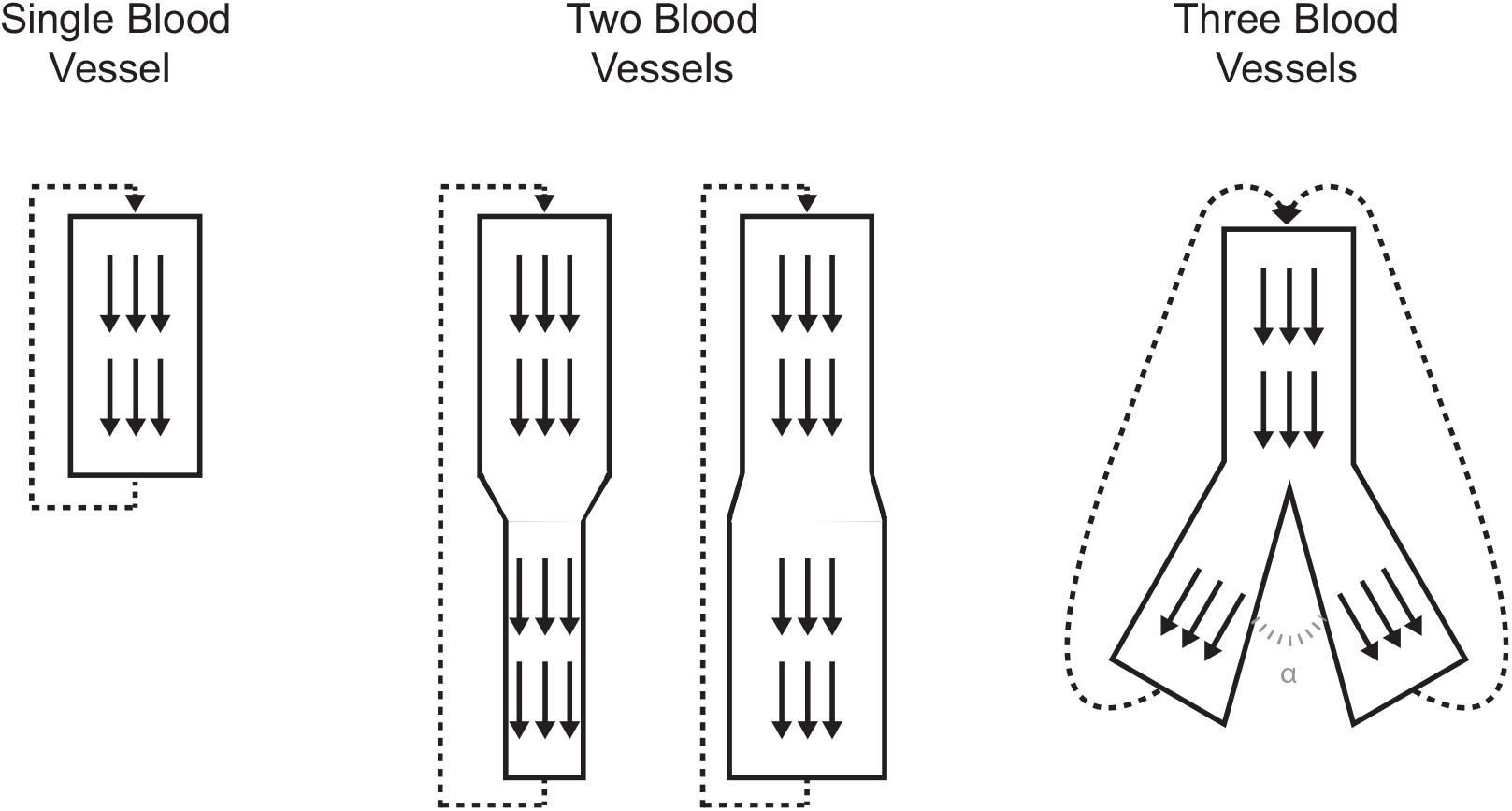
Schematic view of the blood flow in one blood vessel, two blood vessels, and one blood vessel that divides into two with angle *α* between them.

Furthermore, to evaluate the flow dynamics influence for the entire mouse model rather than in the scale of a few blood vessels, we compute the *in silico* results of the model proposed by Lazebnik et al. [46] but replace the flow dynamics (see Eq. (1) in [46]) with the proposed flow dynamics (see Section 4.2.1) and compared with the original model. The comparison was conducted over five *in vivo* experiments as described in Table 2 and evaluate in several points in time, according to the reported *in vivo* values of each experiment.

### 6.2 PKPD dynamics component

To investigate the ability of the model to fit the PKPD dynamics on *in vivo* data, we conducted three tests: a fit of the *in vivo* values for each NP; a fit of the *in vivo* values except the last one which is been predicted, for each NP; and a fit of the *in vivo* values for a group of NPs and prediction of the *in vivo* values of another NP. Therefore, the experiment can be divided into two phases: fitting and predictions. During the fitting phase, the PKPD dynamics parameter (e.i., Eqs. (13-15)) are chosen randomly at first. Given a set of *in vivo* values, the gradient descent algorithm is used to fit the parameters such that the *L*_2_ distance between the *in vivo* values and the *in silico* results over time is used as the error function to minimize. This process is repeated until the change in the error function is smaller than 0.001 (chosen manually). On the other hand, during the prediction phase, the parameters obtained from the fitting phase are used. The model is evaluated on a given set of *in vivo* values. In both the fitting and predicting phases, the *f* function in Eqs. (13-14) is the flow dynamics and computed independently to the PKPD process as part of the flow dynamics calculations.

The model’s fidelity is defined as the slope of a linear regression between the NPs’ distribution in the heart, kidney, liver, lung, and spleen according to the *in silico* tested model and the NPs’ distribution of the *in vivo* experiment [46].

## 7 Results

The results obtained from the experiments are presented as follows. First, we show the influence of the proposed flow dynamics compared to the models proposed in [46, 72] in three levels - single blood vessel flow, one to one blood vessels flow, and one to two blood vessels flow. Second, we show the influence of the proposed PKPD model compared to *in vivo* values in three levels - explanation of the PKPD dynamics using full fit, NP-level partial prediction of *in vivo* values using partial fitting on the same NP, and cross NPs *in vivo* values using full fitting on a set of NPs and full prediction on another NP.

### 7.1 Flow dynamics

The comparison of the proposed flow dynamics with the one proposed by Lazebnik et al. [46] is performed in respect to the flow dynamics *in silico* results of SimVascular [72] due to the lack in *in vivo* data for comparison and as SimVascular shown to be highly biology accurate on small scale sections of the cardiovascular system [72].

A comparison between the proposed flow dynamics with the one proposed by Lazebnik et al. [46] for a single blood vessel is shown in Fig. 5 such that the x-axis is the time in hours from the beginning of the treatment and the y-axis is the simulator’s fidelity compared to SimVascular [72].

**Figure 5:**
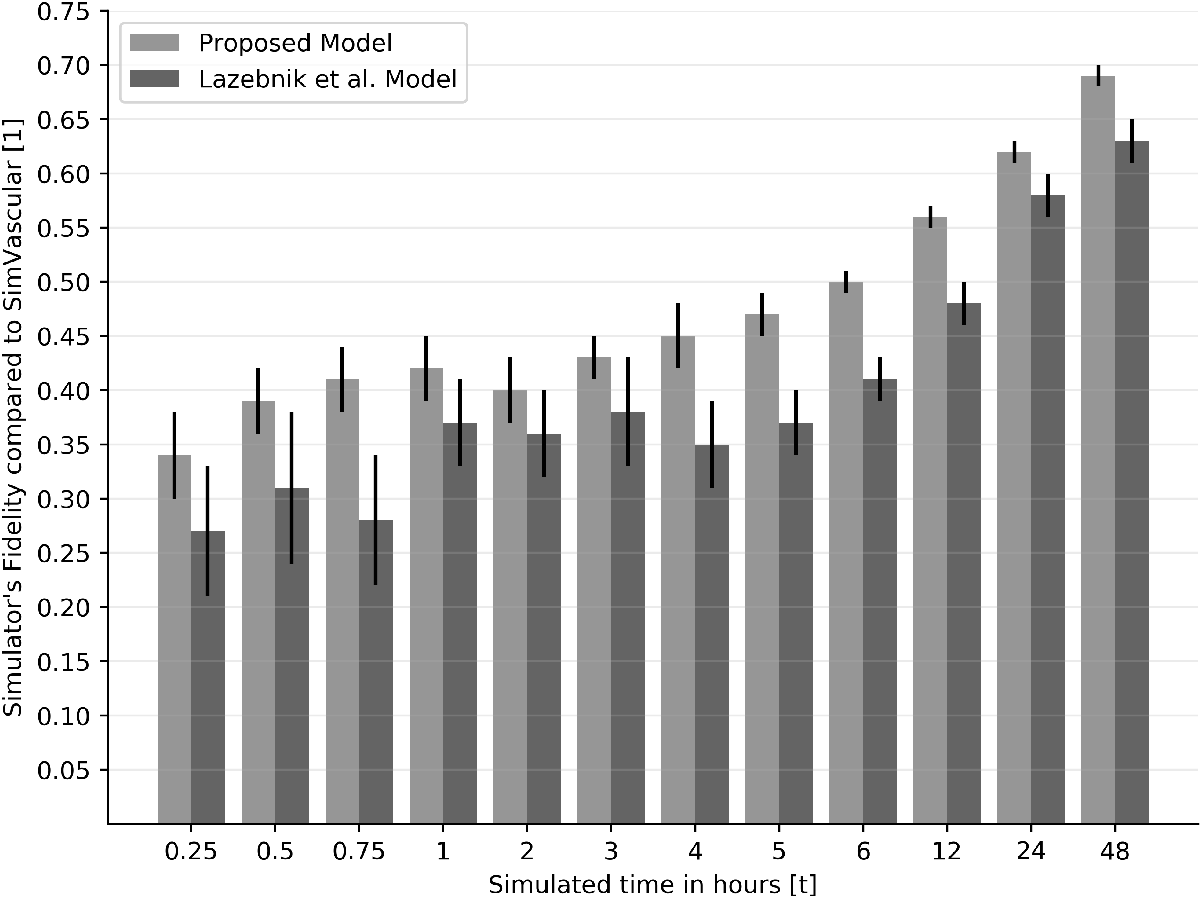
Comparison of the volume of blood flow in a single blood vessel between the model proposed by Lazebnik et al. [46] and the proposed algorithms, in respect to SimVascular [72]. The results are shown as *mean* ± *STD* (standard deviation) for *n* = 5.

Similarly, a comparison between the proposed flow dynamics with the one proposed by Lazebnik et al. [46] for two blood vessels such that the flow is circulating between the first and second blood vessels, as shown in Fig. 6, where the x-axis is the time in hours from the beginning of the treatment and the y-axis is the simulator’s fidelity compared to SimVascular [72].

**Figure 6:**
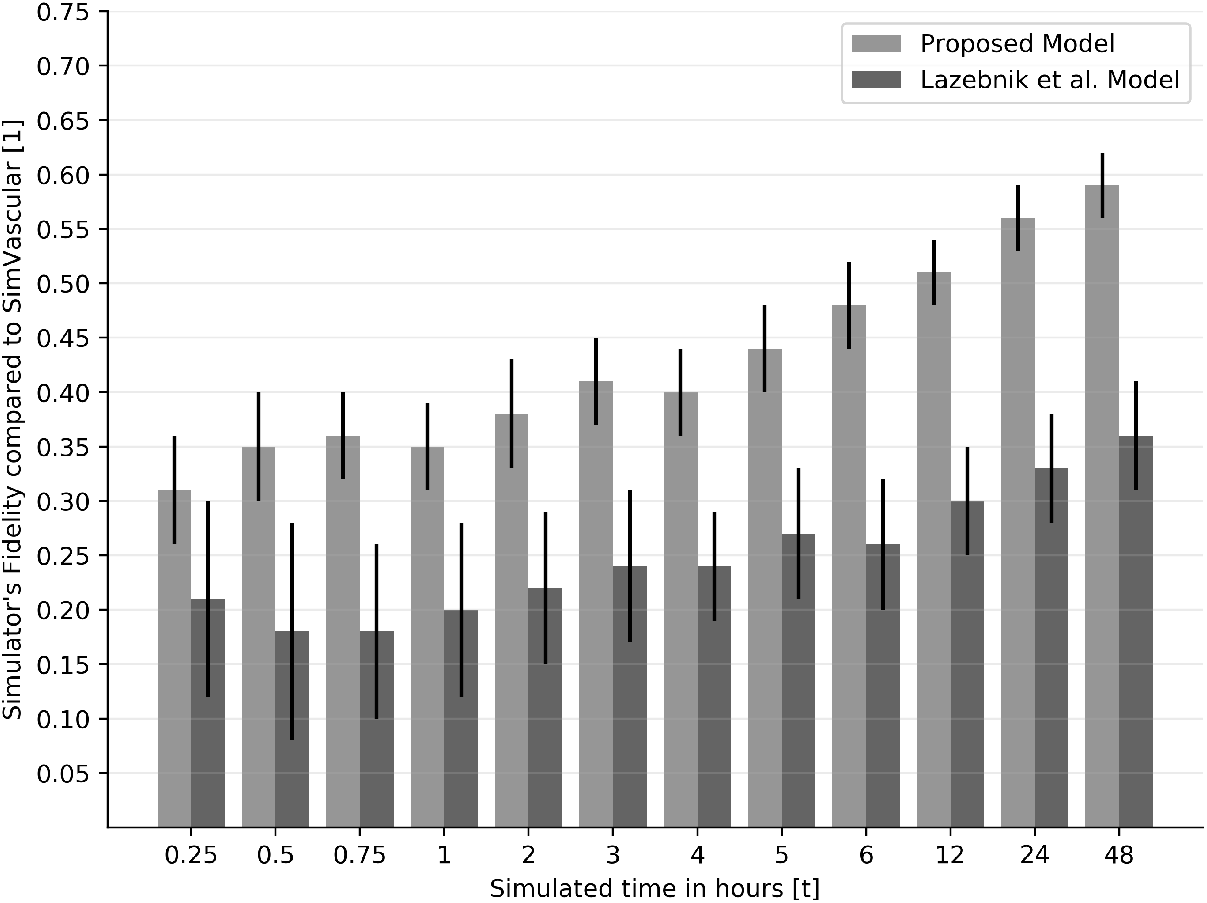
Comparison of the volume of blood flow in two blood vessels between the model proposed by Lazebnik et al. [46] and the proposed algorithms, in respect to SimVascular [72]. The results are shown as *mean* ± *STD* (standard deviation) for *n* = 5.

In addition, to evaluate the influence outer blood vessel geometric configuration, a comparison between the proposed flow dynamics with the one proposed by Lazebnik et al. [46] for three blood vessels such that the flow is circulating between the first blood vessel and both the second and third blood vessels that have an angle *α* between them, as shown in Fig. 7, where the x-axis is the time in hours from the beginning of the treatment and the y-axis is the simulator’s fidelity compared to SimVascular.

**Figure 7:**
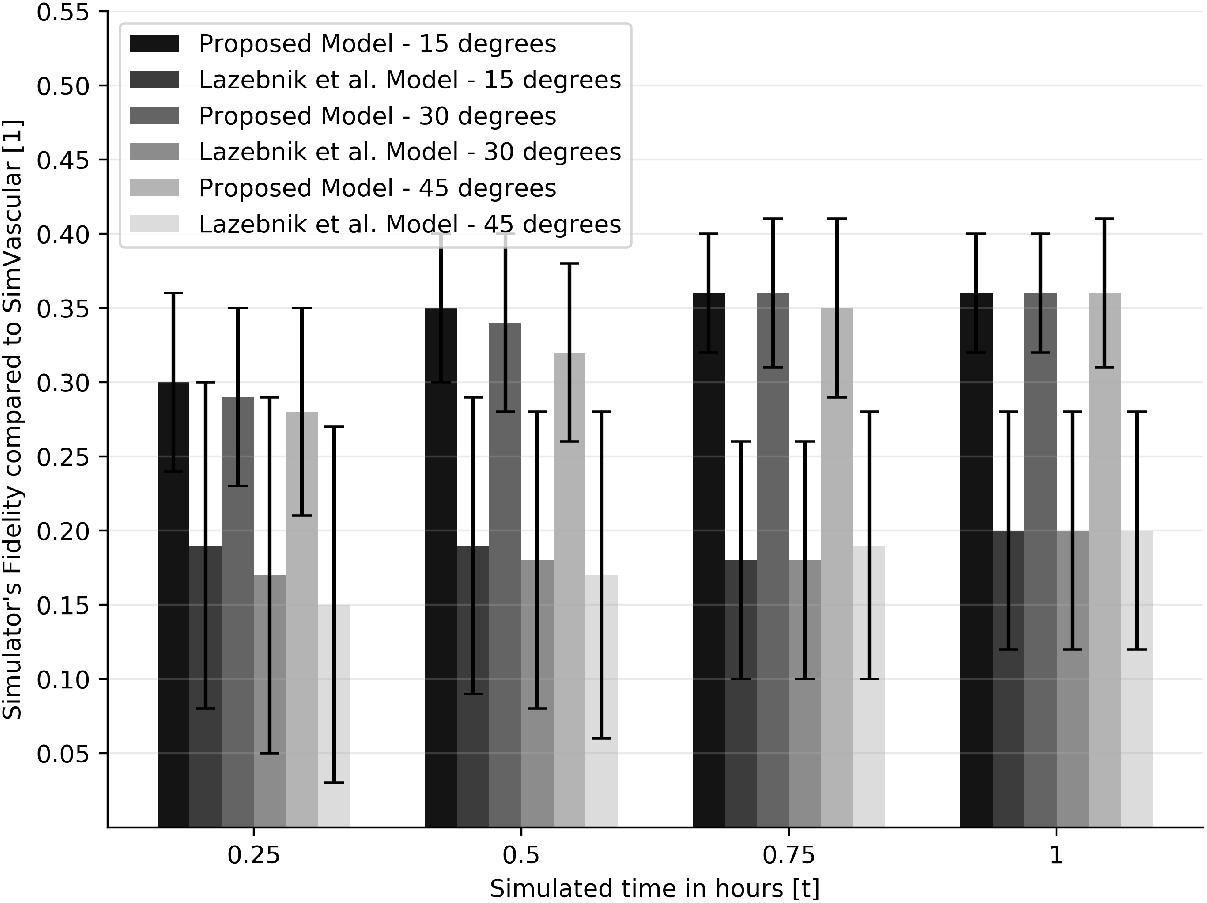
Comparison of the volume of blood flow from a single blood vessel into two blood vessels with an angle of *α* = (15, 30, 45) degrees between them, between SimVascular [72] and the proposed algorithms. The results are shown as *mean* ± *STD* (standard deviation) for *n* = 5 blood vessels.

Furthermore, the evaluation of the flow dynamics in the whole mouse model is shown in Table 3. The table is divided into five *in vivo* experiments, the number of NP types in each experiment, the time in minutes from the beginning of the experiment, and provide a comparison between the model proposed by Lazebnik et al. [46] and the same model but with the proposed flow dynamics in the form of the model’s fidelity. The results are shown as the mean of *n* = 70 repetitions and the mean of all the NPs in each *in vivo* experiment.

**Table 3:**
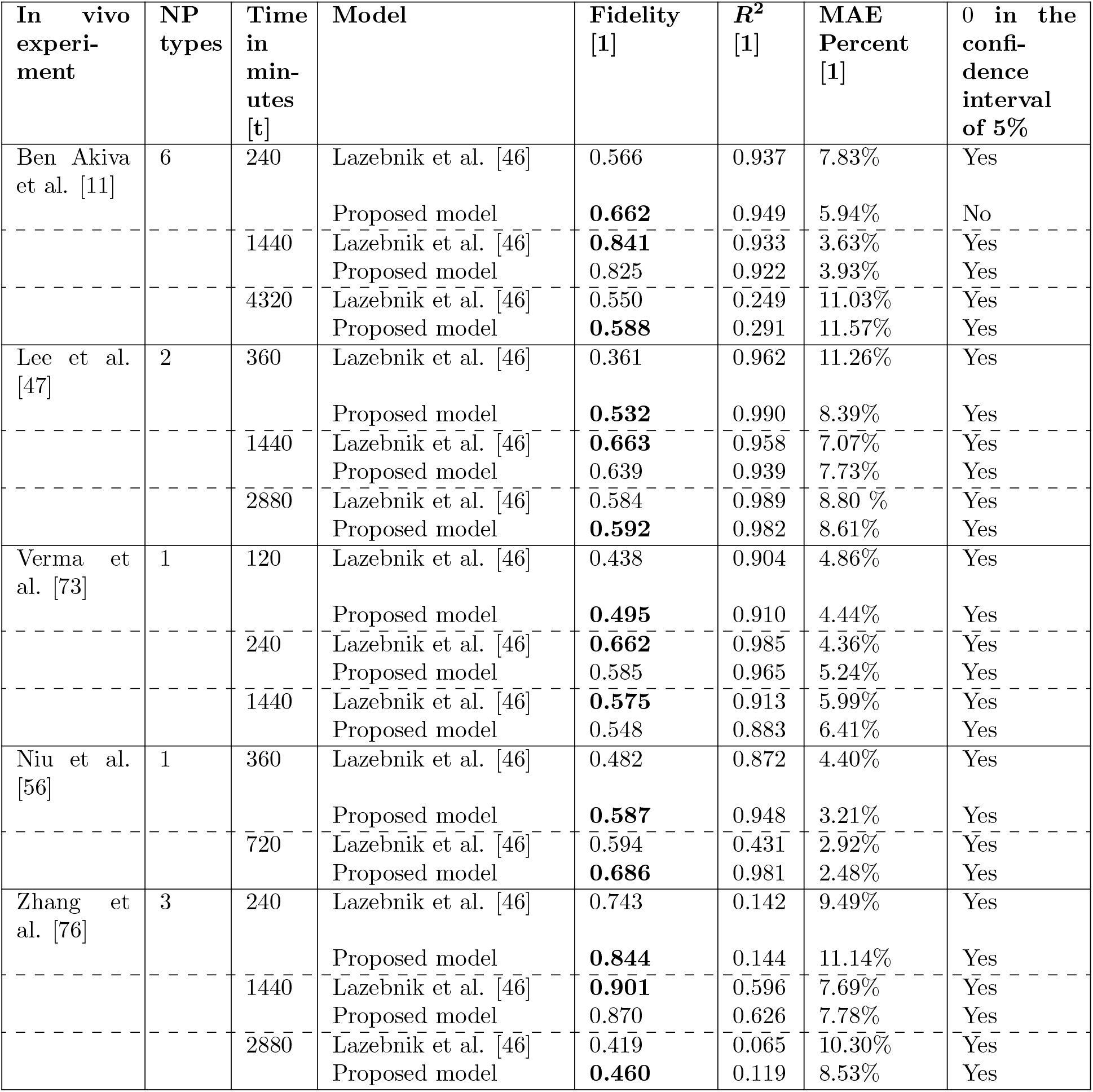
A comparison between the flow dynamics algorithm in [46] and the proposed algorithm on five different *in vivo* experiments. The results are shown as the mean of *n* = 70 repetitions and the mean of all the NPs in each *in vivo* experiment. The confidence interval is obtained by performing a two-tailed T-test between the *in vivo* values (repeated for *n* times) and the *in silico* results.

### 7.2 PKPD dynamics

A full fit of the proposed PKPD model for five organs (heart, liver, kidney, lung, and spleen) for each *in vivo* experiment - divided into four model types: the Lazebnik et al. [46] (namely, the baseline model), improved flow dynamics on top of the baseline model, improved PKPD on top of the baseline model, and the improved PKPD and flow dynamics on top of the baseline model (namely, the combined model), is shown in Fig. 8, such that the x-axis is the time in hours from the beginning of the experiment and the y-axis is the simulator’s fidelity such that the results are shown as a *mean* ± *STD* (standard deviation) of each *in vivo* experiment over *n* = 70 repetitions.

**Figure 8:**
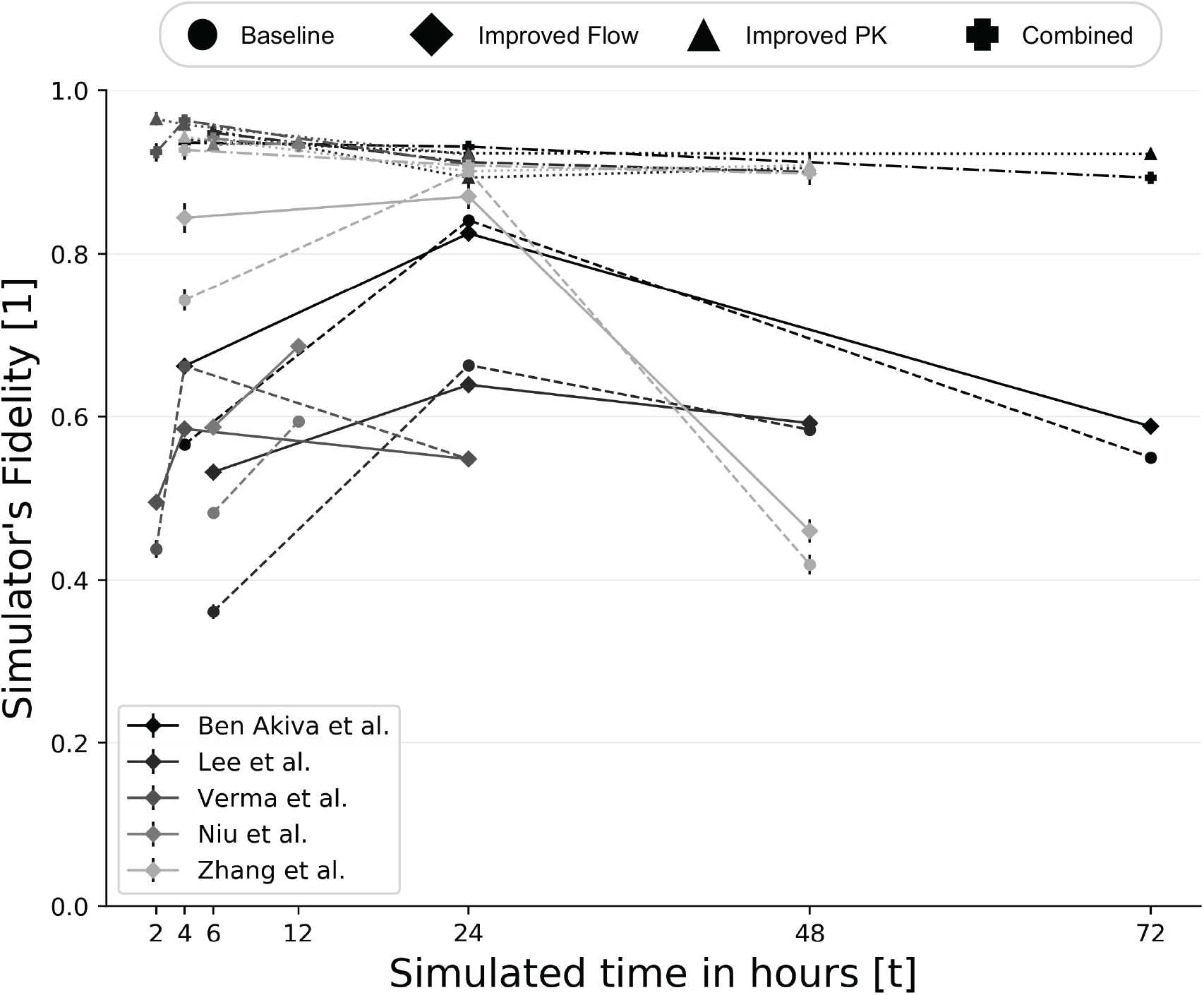
The simulator’s fidelity in respect to the *in vivo* values over time - by fitting the model on all the data. The lines’ colors indicate the *in vivo* experiment has been fitted. The *baseline* model is the model proposed by [46], the *improved flow* model uses the flow dynamics presented in Section 4.2.1, the *improved PK* model uses the PK dynamics presented in Section 5, and the *Combined* model uses both the flow and PK dynamics. The results are shown as *mean* ± *STD* (standard deviation) for *n* = 70.

Differently, Fig. 9 presents the mean simulator’s fidelity for each *in vivo* experiment and model type such that the first two (or one for Niu et al. [56]) *in vivo* values as distribution of the NPs between the organs are used for fitting the model and the last (in time) *in vivo* values are used to evaluate the simulator’s fidelity. In Fig. 9, the x-axis is the *in vivo* experiment and the y-axis is the simulator’s fidelity. The horizontal lines are the weighted (by the number of hours) average of each model type.

**Figure 9:**
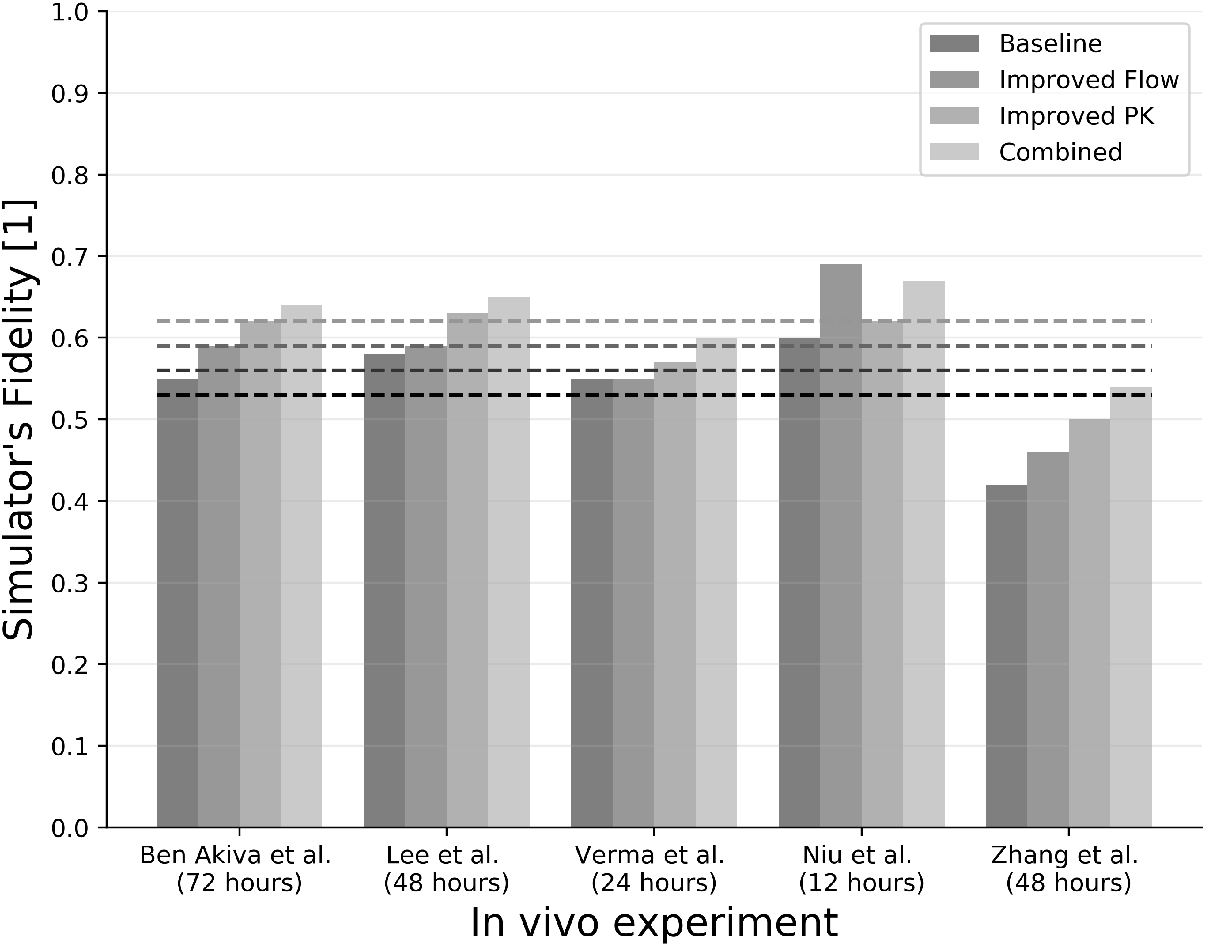
The simulator’s fidelity in respect to the *in vivo* values of five experiments, divided by model’s variations - by fitting the model on each NP (for the first two timestamps) and taking the average for each experiment of a predicted third timestamp. The horizontal lines are the weighted (by the number of hours) average of each model type.

In addition, a cross-NPs prediction such that the PKPD dynamics are fitted to a given set of NPs (Λ) and predicted on another NP *np*_*p*_ such that *np*_*p*_ ∉ Λ from the save *in vivo* experiment is as follows:

- A cross-NPs between the NPs investigated by Ben Akiva et al. [11]. We take one of NP out of the group of six each time (ordered as shown in Table 2) and obtain *v* = [0.52, 0.53, 0.53, 0.56, 0.54, 0.51] fidelity (*E*(*v*) = 0.531, *SD* = 0.016).
- A cross-NPs between the NPs by Ben Akiva et al. [11] that are PLGA coated and sized 200 and 240 nm (see NPs number 1 and 3 in Table 2), respectively was used to predict the 220nm in size and PLGA coated NP from Ben Akiva et al. [11] (see NPs number 2 in Table 2) and obtained as 0.57 fidelity.
- A cross-NPs between the two NPs investigated by Lee et al. [47], resulted in 0.61 fidelity.
- A cross-NPs between the two NPs in size 241nm investigated by Zhang et al. [76], resulted in 0.48 fidelity.

Moreover, the spherical-shaped, 200nm in size, and with PLGA surface NP from Ben Akiva et al. [11] is used as the training data to cross-NPs predict the spherical-shaped, 220nm in size, and PLGA surface NP from Niu et al. [56] and obtained 0.42 fidelity.

## 8 Discussion

This study presents a novel PKPD mathematical model for nanomedicine that takes into consideration two spatio-temporal components: flow dynamics that is based on a cylinder approximation for blood vessels’ geometry with 2-dimensional single-phase Newtonian fluid NS equations, as shown in Sections 4.1. In addition, heartbeat and elastic collision between NPs are taken into consideration in the flow dynamics, as shown in Sections 4.2; and generic PKPD dynamics that take into consideration the natural decay of a drug in the blood-stream and each organ, the decay due to interaction with the immune system and the movement between the organs and the bloodstream. In addition, the immune system’s response due to the stimulus of the drug is taken into consideration in the model, as shown in Section 5. Therefore, the proposed model provides a platform for single-type NPs’ swarm.

The first improvement of the proposed model is its NPs flow dynamics in the cardiovascular system. The proposed approach is a combination between a pure graph-based approach where NPs flow between blood vessels in a directed walk and a fully biophysical numerical simulation that takes into consideration a large number of parameters (that are not necessarily available). The graph-based approach is numerically stable over time but less accurate than the accurate numerical approach, on the other hand, it becomes unstable over time and therefore diverges. The proposed flow dynamics combine the two by relaxing multiple biophysical properties on the one hand and keeping the graph-based approach by treating each vessel separately and the connection between them (corresponding to edges in the graph model).

We evaluated the flow dynamics for small-scale blood flow dynamics with one, two, and three blood vessels compared to the performance of graph-based flow dynamics proposed by Lazebnik et al. [46] and advanced numerical simulation-based flow dynamics provided by SimVascular [72]. For single blood vessel blood flow, the proposed model better approximates the blood flow (0.68 ± 0.02 simulation fidelity compared to SimVascular for 48 simulated hours) but does not statistically significantly improve the results, as shown in Fig. 5. In the case of two blood vessels, the proposed flow dynamics improve significantly - 0.59 compared to 0.35 simulator’s fidelity, after 48 simulated hours. In a similar manner, for three blood vessels, the proposed flow dynamics obtain 0.35 fidelity compared to 0.2 obtained by the model proposed by Lazebnik et al. [46].

When evaluating the model on the mouse model that has 192 blood vessels [46], the proposed model (*Improved flow* outperforms the model proposed by Lazebnik et al. (*Baseline*) for five *in vivo* experiments in the range of 2 to 12 simulated hours. This range can be explained as the flow dynamics are more significant when the NPs do not enter the organs or are evacuated from the bloodstream. Afterward, the difference is mostly dependent on the PKPD dynamics that happen in the organs and indeed the delta between the models is reduced between 12 and 72 simulated hours.

We evaluated the proposed model using five *in vivo* experiments by Ben-akiva et al. [11], Lee et al. [47], Verma et al. [73], Niu et al. [56], and Zhang et al. [76]. For each *in vivo* experiment, there are three point in time (except of Niu et al. [56] that has only two points) in which the distribution of the NPs population over five organs (heart, kidney, liver, lung, and spleen) is obtained. First, we performed a model fitting for each NPs separately and showed the average simulator’s fidelity for each *in vivo* experiment in Fig. 8. For the case where the full model is evaluated (e.i., the “combined” case), the simulator’s fidelity is 0.923 on average with a standard deviation of 0.007. As a result, it is safe to claim that the model well describes the biological processes of NPs-based drug distribution *in vivo*. In addition, we used the first (in time) two distributions of NPs in the organs for each NP separately (one distribution in the case of Niu et al. [56]) to fit the model and the last distributions of NPs in the organs are predicted by the model and compared to the *in vivo* values, as shown in Fig. 9. We show 0.09 improvement in the simulator’s fidelity on average in the proposed model compared to the model proposed by Lazebnik et al. [46] (e.g., the baseline model) by obtaining 0.62 compared to 0.53. Furthermore, the standard deviation of the proposed model is 0.046 which means the model is stable across multiple NPs types (13 samples) with a wide range of properties (such as size, geometry, and surface material).

Furthermore, we used the whole data of each NP on a set of NPs with at least one common property (size, geometry, or surface material) and evaluated the simulator’s fidelity on an additional NP with the same common property. For the case of Ben Akiva et al. [11], the average simulator’s fidelity is obtained to be 0.53 and the standard deviation is 0.02. For comparison, the average simulator’s fidelity prediction with the first two distributions of an NP is used as fitting data, the baseline model in the case of Ben Akiva et al.’s *in vivo* experiment is 0.55; which is only better in 0.02. That means the model can generalize data between NPs. On a larger scale with NPs from different *in vivo* experiments, the average simulator’s fidelity was obtained to be 0.515 and the standard deviation of 0.052. We can therefore conclude it fairly generalizes the PKPD dynamics of different NPs from historical data of other NPs’ *in vivo* experiments but the results are sensitive to the similarity between the fitted data and the tested one.

The proposed model is able to reduce the amount of NPs-based drug biodistribution *in vivo* experiments by estimating the outcomes of such experiment of a new NP-based drug based on similarity with the previous group of *in vivo* experiments and running an *in silico* experiment. In a similar manner, by fitting the model short-term *in vivo* experiment, using the *in silico* model, researchers would be able to approximate the dynamics of the drug on long-term usage.

A possible future work is to obtain the simulator’s results on a large number of NPs-based drugs *in vivo* experiments with a wide range of properties’ values and use this data to train a machine learning model to further improve the generality capabilities of the proposed model, further improving its performance.

